# Comparative study of the effect of neutrons emitted from neutron source ^241^Am-Be and curcumin on MCF-7 breast cancer cells in 3D culture medium

**DOI:** 10.1101/2022.01.25.477792

**Authors:** Sajedeh zargan, Mehdi salehi borough, Jamil zargan, Mohsen shayesteh, Ashkan Haji Noor Mohammadi, Mohsen Mousavi, Hani Keshavarz Alikhani

**Author notes:** Corresponding Author: Mehdi salehi borough, Department of Medical Radiation Engineering, Central Tehran Branch, Islamic Azad University, Tehran, Iran.

## Abstract

**Introduction:** Cancer is one of the major medical problems threatening human health. Breast cancer is the most prevalent type of cancer in women. Reports indicate that treatments such as surgery, chemotherapy, biotherapy, and radiotherapy play a limited but important role in treating the disease. For more efficient treatment, new treatment strategies have been evolved based on combination therapy in which two or more different methods are exploited for this aim. In the present study, a combination therapy based on radiotherapy (using neutron radiation emitted from a ^241^Am-Be neutron source) and biotherapy (using curcumin) is applied to investigate the treatment efficiency of MCF-7 breast cancer in a three-dimensional culture medium.

**materials and methods:** The neutron dose rate in the ^241^Am-Be source was measured using BF3 detection and MCNPX simulation methods, and then the results of both methods were compared. MTT, neutral red uptake assay, nitric oxide, glutathione assay, catalase, cytochrome c, comet assay, and caspase-3 were used to determine the effect and type of mortality due to neutron effect as well as the combined effect of neutron and curcumin in cancer cells.

**Results:** Calculating the fast neutron flux around the source with two methods of simulation with MCNPX and detection with BF3 showed that with increasing distance from the source, the fast neutron flux decreased exponentially. The results of neutron dose rate measurement in the ^241^Am-Be source also showed that if the test cells in the vial are located at a distance of 22 cm from the inlet of the collimator and are exposed to neutron radiation for 5, 10, 15, and 20 hours, the neutron dose received by breast cancer cells will be 3, 6, 9, and 12 mGy/h, respectively.

The results of cytotoxicity due to neutron irradiation as well as the combined effect of neutron and curcumin (80 μM concentration) showed that neutron irradiation with or without curcumin at 5, 10, 15, and 20 hours reduced the survival of tumor cells. According to the results, the rate of apoptosis due to the neutron effect at irradiation times has increased with increasing time. Statistical analysis also showed that the rate of apoptosis due to the combined effect of neutrons and curcumin was not significant compared to the effect of neutrons only. The results of studying the effect of neutrons and the combination of neutrons and curcumin on the production of nitric oxide, catalase, and GSH also showed that curcumin has an antioxidant effect by reducing the amount of nitric oxide and increasing the production of catalase and glutathione in cells. However, neutrons, alone, lacked this effect.

**Conclusion:** Overall, this study showed that the neutron source ^241^Am-Be with the applied doses was able to destroy the c–o, and c–h bonds of curcumin, resulting in cell death and apoptosis induction in breast cancer cells. It was caused by neutron radiation. On the other hand, according to the results of the comet assay and caspase-3 experiments, although neutrons induced apoptosis in breast cancer cells, the death rate due to necrosis was much higher than apoptosis. Due to the significant anti-cancer effect of curcumin in 3D culture, the use of this molecule before or after neutron therapy is recommended.

**Highlights:**

- The results of neutron dose rate measurement in the ^241^Am-Be source showed that if the test cells in the vial are located at a distance of 22 cm from the inlet of the collimator and are exposed to neutron radiation for 5, 10, 15, and 20 hours, the neutron dose received by breast cancer cells will be 3, 6, 9, and 12 mGy/h, respectively.
- The results of cytotoxicity due to neutron irradiation as well as the combined effect of neutron and curcumin (80 μM concentration) showed that neutron irradiation with or without curcumin at 5, 10, 15, and 20 hours reduced the survival of tumor cells.
- According to the results, the rate of apoptosis due to the neutron effect at irradiation times has increased with increasing time. The results of studying the effect of neutrons and the combination of neutrons and curcumin on the production of nitric oxide, catalase, and GSH also showed that curcumin has an antioxidant effect by reducing the amount of nitric oxide and increasing the production of catalase and glutathione in cells. However, neutrons, alone, lacked this effect.
- this study showed that the neutron source ^241^Am-Be with the applied doses was able to destroy the c–o, and c–h bonds of curcumin, resulting in cell death and apoptosis induction in breast cancer cells. It was caused by neutron radiation.
- according to the results of the comet assay and caspase-3 experiments, although neutrons induced apoptosis in breast cancer cells, the death rate due to necrosis was much higher than apoptosis.

## 1.1 Introduction

With the increasing use of radiation in medicine, industry, and research over the past few decades, the need for more radiation protection in radiologists is important (1). Studies show the possibility of further spread of possible late-stage radiation complications such as cancer, genetic damage, and leukemia, which are the most important complications of ionizing radiation in radiation staff. This issue necessitates the importance of advanced research in the field of radiation protection. On the other hand, the use of radiotherapy in the treatment of cancers, including breast cancer, is considered by many medical centers. It is one of the most common cancers in women and the second most common cancer globally (2). According to statistics published by the World Health Organization (WHO) in 2018, one out of every 8 to 10 women suffers from breast cancer. In addition, data published by the Cancer Research Center of Iran in 1398 show that one in 10 to 15 women is affected with this cancer (3).

Today, the discovery of beneficial therapies and effective drugs with the least side effects in patients is on the agenda of many research centers around the world. Neutron therapy is one of the proposed methods for treating some types of cancer (4). Evidence suggests that neutrons are more efficient at causing losses in hypoxic tumor cells, and the amount of damage caused by them is less dependent on the cell cycle (5). Due to the larger relative biological effect (RBE) of neutrons compared to photons, it is now considered one of the treatment choices for treating slow-growing tumors such as breast cancer. One way to reduce the destructive side effects of neutrons is to reduce the dose received by using combination therapies such as biotherapy and neutron therapy. One of the advantages of this method is the reduction of systemic toxicity caused by radiation therapy due to the decrease of neutron dose received by patients (6). One of the biomolecules with anti-cancer properties proposed for cancer treatment is the curcumin in turmeric, which has extensive biological properties such as anti-inflammatory, antioxidant, anti-diabetic, and anti-cancer activity (7, 8).

In this study, for the first time, the combined effect of neutron radiation produced by ^241^Am-Be and curcumin on breast cancer cells (MCF-7) under three-dimensional cell culture was investigated. Three-dimensional cell culture is one of the best laboratory models used to study the viability of cancer cells due to the effects of chemicals, biomolecules, and radiation by creating natural conditions similar to living organisms (9).

## 1.2. materials and methods

### 1.2.1 Materials and equipment

### 1.2.2. Materials and equipments for the measurement of the neutron dose

^241^Am-Be with 5-curie activity with relevant protection, BF3 detector made by Germany’s centronic company, preamplifier, amplifier, high voltage source, counter, MCNPX code, 11 cm cell culture flask

### 1.2.3. Materials and equipments for investigating the effect of neutrons and curcumin on cancer cell death

MTT- DMSO- Sodium hydroxide- Sydrochloric acid- Ethidium bromide- Antibiotic - Antimycotic- Penicillin- Agarose- Acetic acid- Triton x-100- Phosphoric acid- Flat bottom 24-wells plate- Flat bottom 96-wells plate- Glycerol- Neutral red dye- Coomassie brilliant blue G-250- DMEMF12, DMEM- Trypsin-EDTA -Fetal bovine serum- Phenol red -Trypan blue-Curcumin

In this experimental-laboratory study, breast cancer cells (MCF-7 IBRC C10082) were prepared from the cell bank of Imam Hossein University.

### 1.2.4 Measurement of neutron dose rate in ^241^Am-Be source

The neutron dose rate in the ^241^Am-Be source was measured using BF3 detection and MCNPX simulation methods, and then the results of the two methods were compared. After confirming the results, the absorbed dose of the desired vial/flask containing cancer cells was calculated under the three-dimensional cultures used in the experiments. In this method, to measure the absorbed dose of the cell culture vial/flask, a cylinder with vial/flask dimensions was defined in the space inside the collimator, and the material inside it was considered water. The reason for choosing water is the type of vial/flask used for cell culture and the effect of neutrons and curcumin on it in three-dimensional culture. The flask and tubes used in this study are made of polystyrene with a density of 1gr/cm^3^, approximately equal to the density of water and soft tissues. Absorption dose was measured in two modes with BF3 detector and simulation by MCNPX method at four distances of 20, 24, 28, and 32 cm from the center of the ^241^Am-Be source. Due to the length of the BF3 detector (31 cm), the middle of it was considered as a sensitive point. In measuring the absorption dose by the MCNPX method using F6 Tally, the absorption dose was directly calculated in terms of MeV/gram. Then, the maximum dose rate was calculated by determining the distance of the vial from the inlet of the collimator and measuring the cylindrical absorption dose with the dimensions of the vial/flask of cell culture from the two detection methods, BF3 and MCNPX. The dose rate was measured at a distance of 22 cm from the source at the neutron irradiation times specified for cell experiments.

### 1.2.5 Comparative effect of neutron and curcumin on mortality of breast cancer cells (MCF-7) under 3D culture

#### 1.2.5.1 Encapsulation of cells in alginate

To achieve a 3D environment, we encapsulated the MCF-7 cells in alginate particles. At first, the alginate solution was produced by dissolving 0.12g of sodium alginate powder (Sigma-USA) in 10 mL of 0.9% sodium chloride solution. The alginate solution was filtered by a 0.22 μm plastic syringe filter. 2×10^4^ MCF-7 cells were seeded in each well of 96-wells plate and its volume set to 1mL with alginate solution. Cell individualization and suspension of cells in alginate solution were performed using a 22G plastic syringe. Alginate capsules were produced by injecting a cell-alginate mixture by a 22G plastic syringe into a bath of 100 mM calcium chloride (drops were released into the bath from a distance of 5 cm above the bath’s surface). The capsules were allowed to be polymerized for 10 min. In the next step, after the removal of calcium chloride, the capsules were washed three times with PBS solution (pH 7.4). The washing solution was replaced with 1mL of DMEM containing 10% FBS. The cell capsules were incubated at 37 ° C, with 95% humidity and 5% CO2 (10).

#### 1.2.5.2 Depolymerization of alginate capsules and release of cells

To perform some experiments, such as the alkaline comet test, GSH, catalase, and comet assay in 3D culture following the exposure of cells with different concentrations of curcumin (for 24 h), it was first necessary to depolymerize the cell-containing capsules. The release steps were as follows. The capsules were carefully transferred to 2 mL microtubes. Each microtube was washed thoroughly and several times with PBS. 1 mL of 50 mM sodium citrate solution was added to each microtube, and the tubes were incubated in the laboratory for 20 min. To completely dissolve the capsules and release the cells, microtubes were centrifuged at 1500 rpm for 5 min. The supernatant was discarded, and the cell pellet was suspended in PBS. The microtubes were centrifuged at 1200 rpm for 3 min. The supernatant was discarded, and the sediment containing the cells was used for the mentioned experiments(9).

#### 1.2.5.3 Mixing curcumin in the suitable buffer and removing contamination

Curcumin was obtained with 95% purity from Karen Pharmaceutical Company. 0.02 mg curcumin was dissolved in sterile Dulbecco’s modified Eagle’s medium (DMEM) containing 1% antibiotic-antimycotic and incubated overnight in a CO2 incubator (37 °C and 5% CO2) to remove biological contaminants. The final concentration of curcumin was 80 μM.

#### 1.2.5.4 cell cultures and tests

The results of studies investigating the effect of curcumin on MCF-7 cell growth under three-dimensional culture conditions showed that the IC50 of this substance was 80 μM (11). This concentration was used to evaluate the combined effect of neutron radiation and curcumin on cancer cell growth. In this study, cells encapsulated in alginate were incubated in a flask containing culture medium in two states with curcumin and without it at a distance of 22 cm from the ^241^Am-Be source for 5, 10, 15, and 20 h. Based on the results obtained in this study, these cells were exposed to fast neutrons at 3, 6, 9, and 12 mGy/h, respectively, at the mentioned times. After this time, the flasks containing the cells were transferred to an incubator. The total duration of cell contact with neutrons and curcumin and storage in the incubator was 24 h. The effect of neutrons on breast cancer cell mortality in the presence and absence of curcumin under 3D culture in the above conditions was performed by MTT, neutral red, comet assay, cytochrome c, nitric oxide, catalase,caspase-3, and glutathione assays.

#### 1.2.5.5 Determination of cell viability via MTT assay)

This method was performed according to the ones conducted by Mossman et al. (12). Alginate capsules (containing approximately 1×10^4^ cells) floating in an 11 cm flask containing culture medium treated with 80 μM of curcumin and exposed to neutron radiation at a distance of 22 cm from the ^241^Am-Be source. An equivalent flask without curcumin was used as a control. Samples were incubated for 5, 10, 15, and 20 h. Then, the flasks were transferred to an incubator (5% CO2, 80% humidity, and 37 °C. The total duration of cell contact with neutrons and curcumin and storage in the incubator was 24 h. After that, alginate capsules were transferred to a 96-well plate (three replicates). 20 μL of MTT solution (5 mg/mL) was added to each well. The plate was transferred to a CO_2_ incubator till dark blue crystals (Formazon) were formed. The contents of each well were removed, and after washing with PBS, 200 μL DMSO was added to each well. To completely dissolve the precipitate, the plate was incubated for 2–4 h in the dark condition at room temperature. In this test, a culture medium was used as a control. Indeed, a culture medium containing cells was used as another control. The light absorption of the wells was measured at 570 nm by a microplate reader (Biotek, USA). This experiment was repeated three times, and three wells (3 times) were considered for each concentration. Cell viability was calculated by the following formula (13).

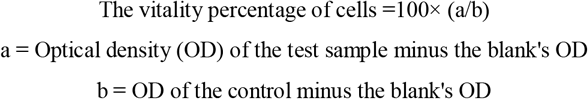

#### 1.2.5.6 Neutral red (NR) uptake assay

NR uptake assay was used to verify the MTT assay results (Winckler 1974) (14). The Neutral red test is based on the ability of live cells to combine and link neutral red to lysosomes (12). The steps of the neutral red colorimetric test are similar to MTT assay test, except that instead of adding the MTT solution to the cell capsules transferred to the microplate, 4 μL of neutral red dye (5 μg/mL) was added to each well and incubated for 1 h at 37 °C, 5% CO2 and 80% humidity under dark conditions, until red crystals formed. Then, the solution in each well was discarded and washed twice with PBS. After that, 200 μL of fixing buffer (formaldehyde 37% (v/v), CaCl2 (10% (w/v) in water) was added to each well and incubated for 1 min. After 1 min, 100 μL of solubilizing buffer (acetic acid0.5%) was added, the plate was incubated in a shaker for 20 min in the dark condition, and the absorbance was measured at 540 nm by a microplate reader (Biotech, USA). Inhibitory percentage due to the combined effect of neutrons and curcumin on cell growth was calculated using the following formula.

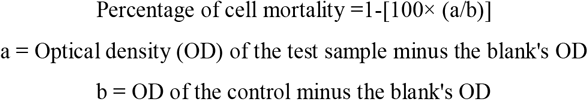

#### 1.2.5.7 Nitric oxide (NO) assay

This method was performed according to Zargan et al. (16-15). The steps of the NO assay were similar to the MTT assay, except that instead of adding the MTT solution to the microplate-transferred cell capsules, 100 μL of the media were transferred to a 96-wells plate and mixed with the equal volume of Griess reagent (Sigma, USA) (0.04 g/mL in PBS, pH 7.4) and incubated for 10 min at room temperature. Absorbance at 520 to 550 nm was measured by a microplate reader (Biotech, USA). Nitric oxide concentration (μM/mL) in treated cells was calculated using sodium nitrite standard curve.

#### 1.2.5.8 Reduced glutathione (GSH) assay

This method was performed according to Sedlak and Linsay (17). The steps of the GSH test were similar to the NO test, except that encapsulated cells containing 5×10^5^ cells were transferred to 1.5 mL tubes for testing. 200 μL of the lysis buffer was added to each well, and protein concentration was assessed by Bradford assay. 40 μL of the obtained solution was removed and transferred to new tubes. 40 μL of 10% TCA solution was added to each tube, and the plate was stored at 4 °C for 2 h. The centrifugation was performed at a speed of 1500 rpm for 15 min. The supernatant was transferred into a clean tube. Then, 75 μL of lysis buffer, 55 μL of Tris HCl buffer (pH=8.5), and 25 μL of DNTB was added to 20 μL of the supernatant. OD of the samples was measured at 412 nm.

#### 1.2.5.9 Catalase enzyme activity assay

This method was performed according to the method described by Sinha et al. (18). The steps of the catalase test in this study were similar to the GSH test, except that after protein measurement by the Bradford method, 5 μL of samples were mixed with 50 μL of the lysis buffer (20 μL DDW and 25 μL of 15% H2O2). Samples were incubated for 2 min at 37 °C and were mixed with 100 μL of potassium dichromate solution. At this stage, the soluble pink color with the blue color of the upper part of the solution was visible. Next, samples were incubated at 100 ° C for 10 to 15 min until the green color appeared. The tubes were spun, and 150 μL of the solution in each well was transferred to a 96-well plate, and the OD was measured at 570 nm using a plate reader (Biotech, USA).

#### 1.2.5.10 Cytochrome c assay

The steps of the cytochrome c test were similar to the GSH test, except that capsules containing 1 ×10^6^ cells were transferred to 1.5 mL tubes for testing. The cells were dissolved in 1 mL of cytosolic purification buffer and incubated on ice for 10 min and homogenized in a Dounce tissue grinder. The homogenized samples were transferred to a new 1.5 mL tube and centrifuged at 700 g for 10 min at 4 ° C. The supernatant was transferred to new 1.5 mL tubes and centrifuged at 10,000 g for 30 min at 4 ° C. The soluble solution, which was the cytosolic component, was collected. Protein concentration was measured using Bradford, and OD was measured at 495 nm.

#### 1.2.5.11 Alkaline comet assay

This method was performed according to the procedure performed by Zimmermann et al. (19). The initial steps of the alkaline comet test were similar to the MTT method, except that after receiving the radiation and incubating in the incubator, the capsules containing 12 × 10^4^ cells were transferred to 1.5 mL tubes three times. Then, 200 μL of PBS was added to each tube containing the sediment of the released cells from the dissolved alginate capsules, and cells were singled out using a needle and isolated. Slides were covered by normal melting agarose (1% (v/v)) and incubated for 10 min at 4 °C. Cell suspensions were mixed with low melting agarose (1% (v/v)) (1:2 ratios) and were applied to the slides. To form a cell layer, a coverslip was applied to spread the preparation. To do cell lysis and nucleus distraction, all slides were incubated for 16–18 h in fresh and cold lysis buffer (NaCl 2.5M, EDTA 100 mM, Tris 10 mM, NaOH 0.2 M, and Triton X-100 %1; pH 10) at 4 °C. Then, slides were washed twice with electrophoresis buffer for 20 min at 4 °C and electrophoresed for 45 min at 4 °C (25 V and 300 mA). For neutralization, the slides were incubated for 10 min in the neutralizing buffer (Tris 0.04 M, pH 7.5). Then, the slides were incubated in 100 μL ethidium bromide (20 μg/ mL) for 10 min at room temperature. Slides were washed two times (10 min each) with water and analyzed by an inverted fluorescent microscope (Nikon, Japan), and results were statistically analyzed.

#### 1.2.5.12 Caspase-3 activity test

Caspase-3 activity was evaluated using the commercial Caspase-3/ CPP32 laboratory kit according to the manufacturer’s instructions (12). The steps for testing the activity of caspase-3 were similar to the cytochrome c test, except that after dissolving the alginate capsules, the resulting cell precipitate and cell plate were suspended again in 100 mL of cold lubricating buffer (…) and incubated on ice for 20 min. Samples were evaluated for protein content by ??. The final volume of 50 μg of the protein content of the samples was then increased to 100 μl using a lubricating buffer and combined with a reactive buffer (containing 10 mM DTT). In the next step, 5 μL of DEVD-pNA 4mM (200 µM) was added to each sample and incubated for 1 h at 37 ° C. In the last step, the absorbance of the samples was measured at 405 nm.

## 1.3. Statistical analysis

Comparison of anticancer effect of neutron and curcumin on MCF-7 breast cancer cells was evaluated by GraphPad InStat software and two-way ANOVA. Differences were considered to be significant at *P* < 0.05(*), *P* < 0.01(**), *P* < 0.001(***) and *P* < 0.0001(****). All experiments were conducted three times.

## 1.4. Results

### 1.4.1 Results of neutron dose measurement in the ^241^Am-Be source using BF3 detection and MCNPX simulation methods

The results of measuring the rate of cylindrical absorption dose with vial/flask dimensions of cell culture from BF3 detection and MCNPX simulation methods are presented in Table 1-1. Based on the dose measurements of the cylindrical absorption dose with the dimensions of a vial/flask of cell culture (length 11 cm) with two detection methods mentioned and at a distance of 22 cm, the maximum dose rate was 0.6 mGy / h. Also, the results of neutron dose rate measurement in the ^241^Am-Be source showed that if the test cells in the vial are 22 cm away from the collimator inlet and are exposed to neutron radiation for 5, 10, 15, and 20 hours, the neutron dose Received by breast cancer cells will be 3, 6, 9 and 12 mGy, respectively.

**Table 1-1:** Absorption dose rate from two detection methods with BF3 and simulation by MCNPX method at different distances from 241Am-Be source.

| **Absorption dose simulated by**<br>**MCNPX method (mGy/h)** | **Experimental absorption dose**<br>**of BF3 detector(mGy/h)** | **Distance from the center of the**<br>**source(cm)** |
| --- | --- | --- |
| 0/67 | 0/61 | 20 |
| 0/45 | 0/40 | 24 |
| 0/31 | 0/28 | 28 |
| 0/24 | 0/21 | 32 |

### 1.4.2. Results of the comparative effect of neutron and curcumin on breast cancer cell mortality (MCF-7) in 3D culture

#### 1.4.2.1 MTT Assay

The results showed that the survival rate of MCF-7 cells in the presence of neutron radiation at 5, 10, 15, and 20 h was 90, 65.96, 62.1, and 53.06%, respectively. This effect was 89.7%, 61.3%, 64.9%, and 50.2% in the presence of curcumin with a concentration of 80 μM at the mentioned times (Figure 1-1). The results showed that neutron irradiation had a significant effect on cells’ viability in the presence of curcumin and in the absence of curcumin compared to control except at 5 h (Figure 1-1). Analysis of the results also showed that the effect of neutron radiation in the presence of curcumin and without it at times 5, 10, 15, and 20 h was not significant (*P*≥0.05)

**Figure 1-1.**
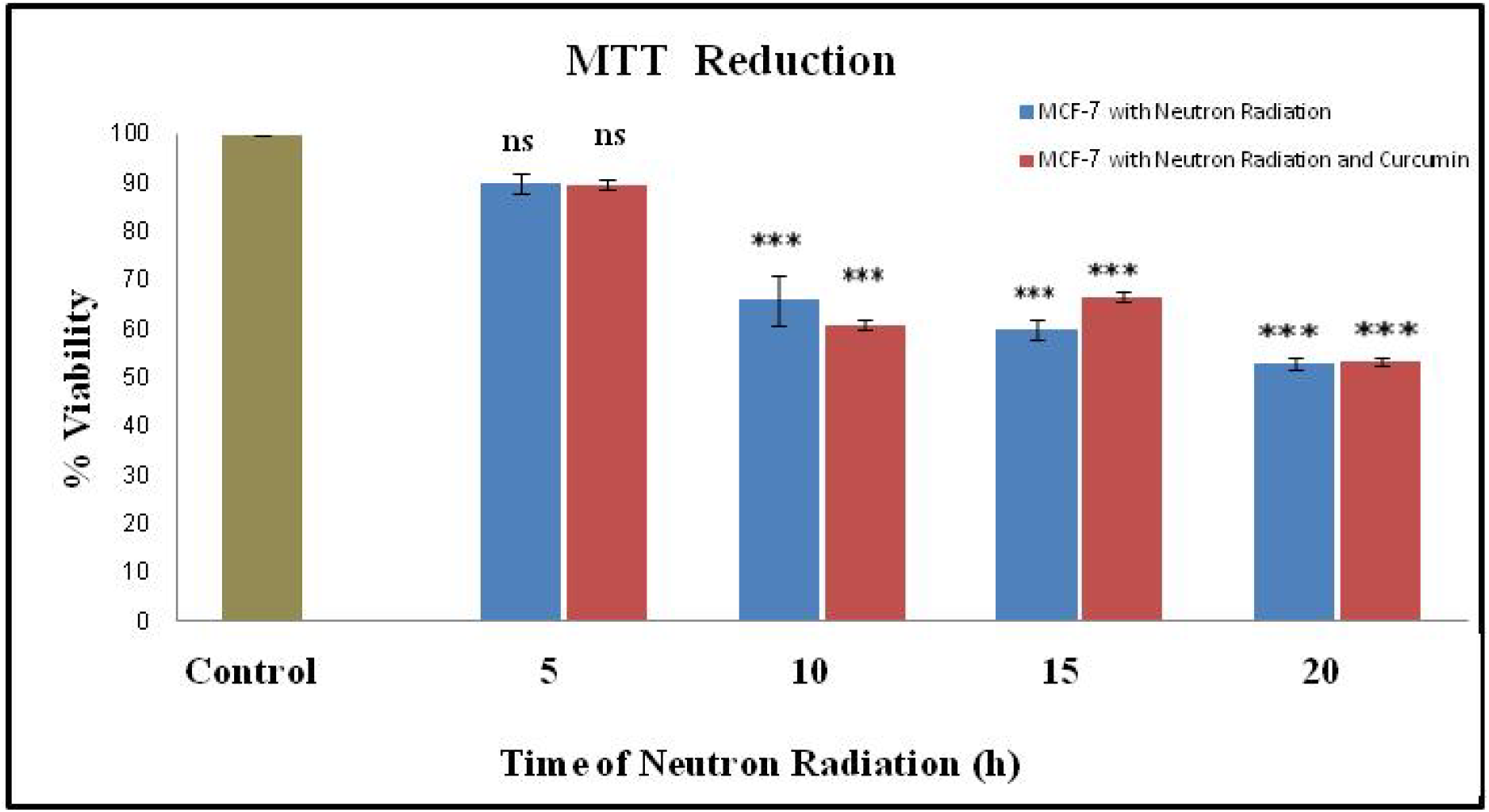
Calculated survival percentage to evaluate the effect of neutrons on breast cancer cell mortality (MCF-7) in the presence and absence of curcumin under 3D culture in the MTT assay. ns: not significant, (***p<0.001)

#### 1.4.2.2 Neutral red uptake assay

The results showed that the percentage of growth inhibition of MCF-7 cells in the presence of neutron radiation at 5, 10, 15, and 20 h was 14.76, 36.63, 30.53, and 22.96, respectively. This rate was equal to 8.76, 30.5, 30.7, and 26.8 in neutron irradiation conditions in the presence of curcumin. Analysis of the results showed (Figure 1-2) that the neutron radiation in the presence and absence of curcumin at all times had a significant inhibitory effect on cells growth. However, the results of neutron radiation at the above times in the presence and absence of curcumin were not significantly different.

**Figure 1-2.**
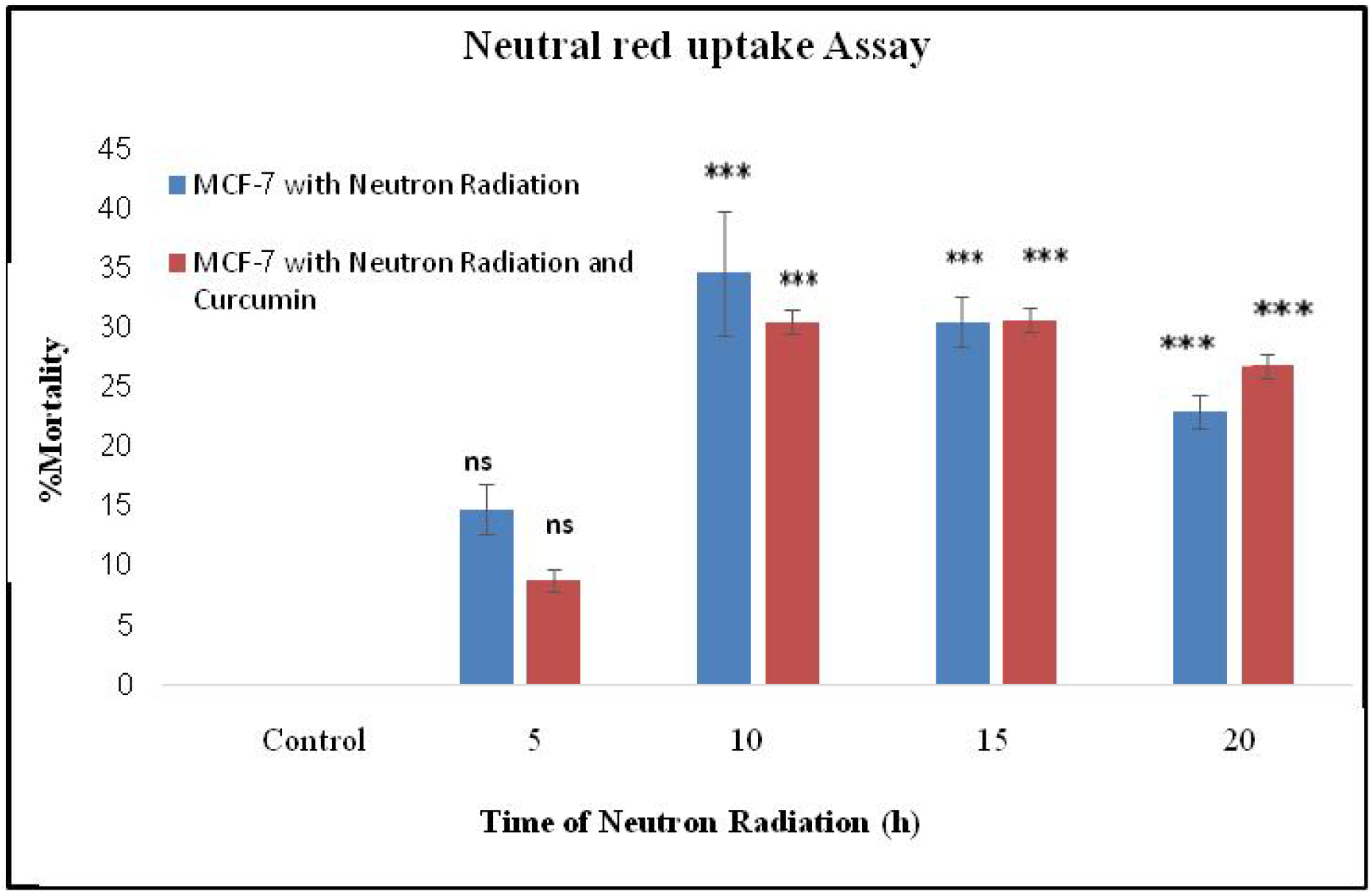
Percentage inhibition of MCF-7 cell growth under neutron irradiation in the presence and absence of curcumin under 3D culture using a neutral red test.(***p<0.001)

#### 1.4.2.3 Nitric oxide assay

The amount of NO release from MCF-7 cells due to neutron irradiation at 5, 10, 15, and 20 h was equal to 2.15, 2.06, 1.83, and 1.78 (Molarities of sodium nitrite). As shown in Figure 1-3, the amount of nitric oxide released from cells by neutron irradiation at all times was not significant (*P*>0.05). The amount of nitric oxide released from cells under neutron irradiation in the presence of curcumin was equal. With 2.55, 2.31, 1.29, and 2.02 that this effect was significantly different in 5 and 10 h with control. The results also showed that the effect of neutron radiation with and without curcumin at times The mentioned were not significantly different from each other.

**Figure 1-3.**
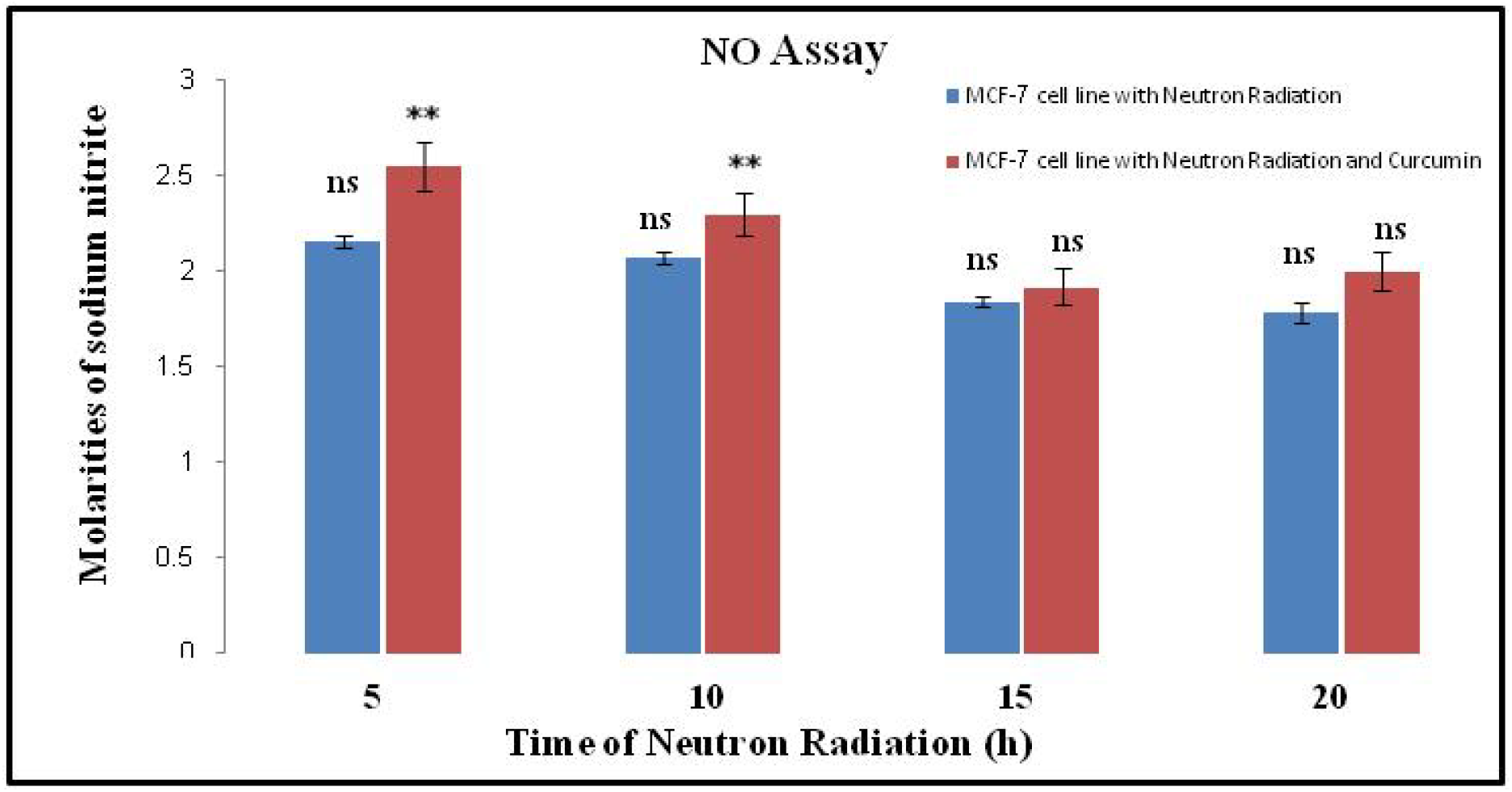
The amount of nitric oxide generated for neutron effect on breast cancer cells (MCF-7) in the presence and absence of curcumin under 3D culture using NO assay. (ns: not significant, **p<0.01).

#### 1.4.2.4 Catalase enzyme activity assay

The results showed that the amount of cellular catalase generated by neutron irradiation at 5, 10, 15, and 20 h was equal to 32.94, 57.43, 32.12, and 33.09, respectively. As shown in Figure 1-4, at all times, except at 10 h, the produced catalase in treated cells was significantly higher than in untreated cells. The produced catalase after the neutron irradiation in the presence of 80 μM curcumin at 5, 10, 15, and 20 h was 38.32, 42.42, 31.03, and 33.77. The produced catalase was not significantly different from the control at all times. The result also showed that the effect of neutron radiation in the presence of curcumin compared to the absence of curcumin was not significantly different, except at 20 h.

**Figure 1-4.**
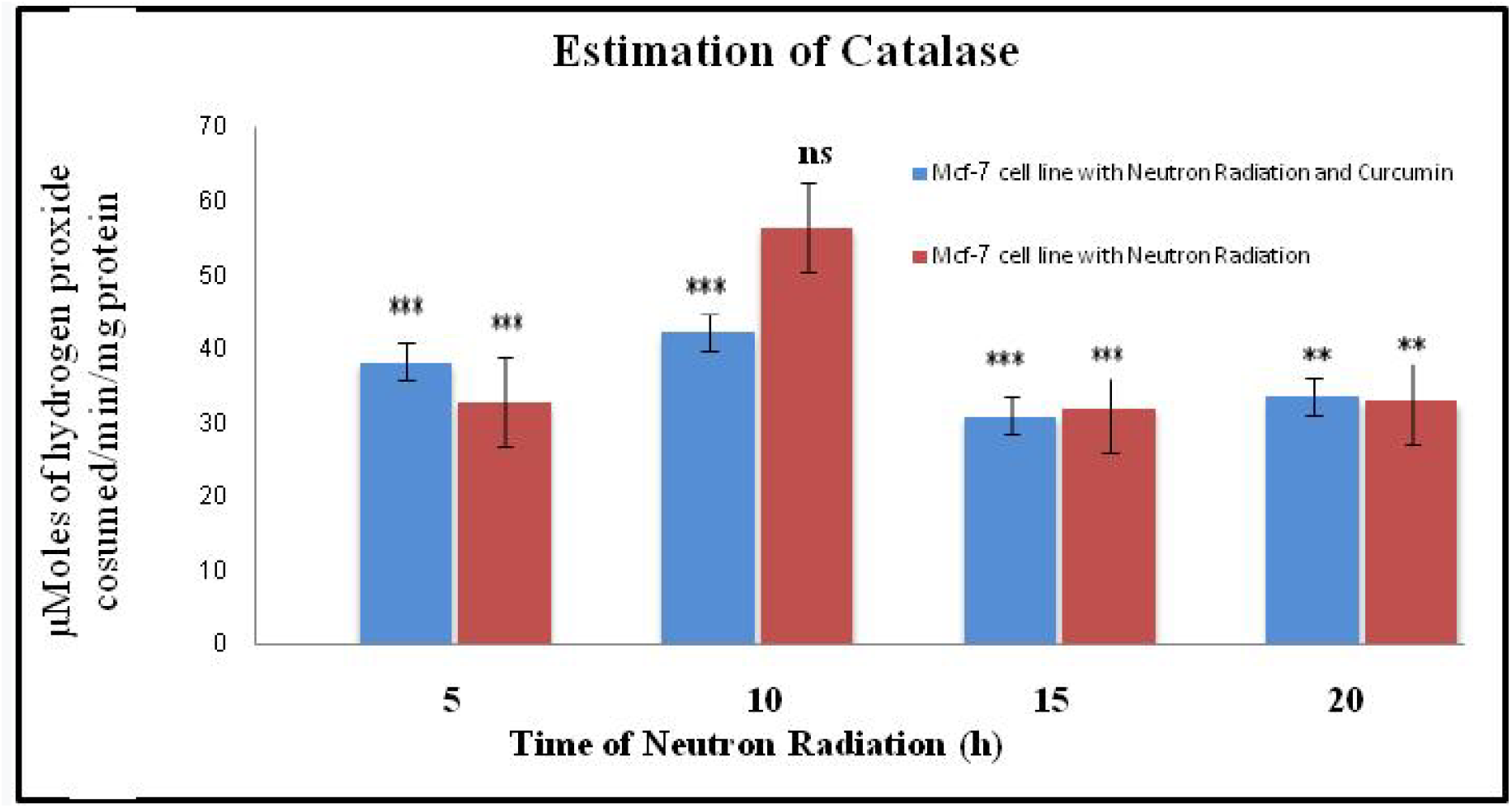
The amount of catalase calculated for the effect of neutrons on breast cancer cells (MCF-7) in the presence and absence of curcumin under 3D culture using the Estimation of catalase test. (ns: not significant, **p<0.01, ***p<0.001)

#### 1.4.2.5 GSH assay

The results showed that the cellular glutathione level produced under neutron radiation at 5, 10, 15, and 20 h was 0.23, 0.40, 0.27, and 0.28 (μg of GSH /mg protein). The amount of glutathione produced after neutron radiation in the presence of curcumin at the above irradiation times was 0.33, 0.31, 0.27, and 0.30 (μg of GSH /mg protein), respectively. As shown in Figure 1-5, neutron radiation in the presence and the absence of curcumin significantly increased the level of cellular glutathione at all times. The presence or absence of curcumin had no significant effect on glutathione production at all times, but at 5 h, is significant.

**Figure 1-5.**
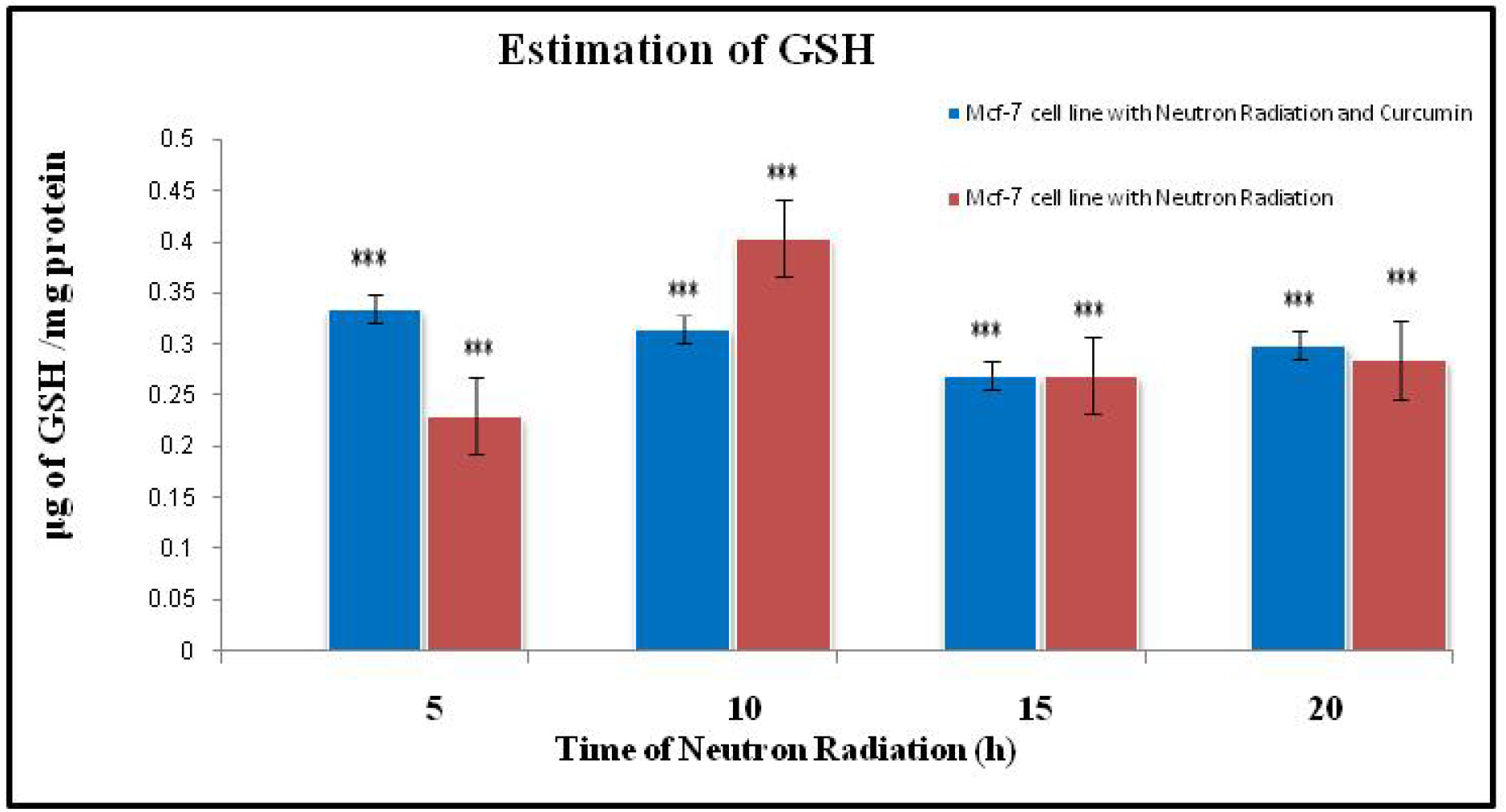
The amount of glutathione calculated by examining the effect of neutrons on breast cancer cell mortality (MCF-7) in the presence and absence of curcumin in 3D culture using the Estimation of GSH assay. (***p<0.001)

#### 1.4.2.6 Cytochrome c assay

The results showed that the amount of cytochrome c released into the cytosol in MCF-7 cells in 3D culture following neutron irradiation at 5, 10, 15, and 20 h was 0.137, 0.130, 0.124, and 0.123. On the other hand, this amount was 0.151, 0.40, 0.123, and 0.124 following the neutron irradiation in the presence of 80 μg curcumin. As shown in Figure 1-6, the effect of neutron radiation in the presence and absence of curcumin had a significant inhibitory effect on cell growth at all times. The results also showed that the presence or absence of the curcumin in combination with the neutron radiation had no significant effect on cell growth inhibition.

**Figure 1-6.**
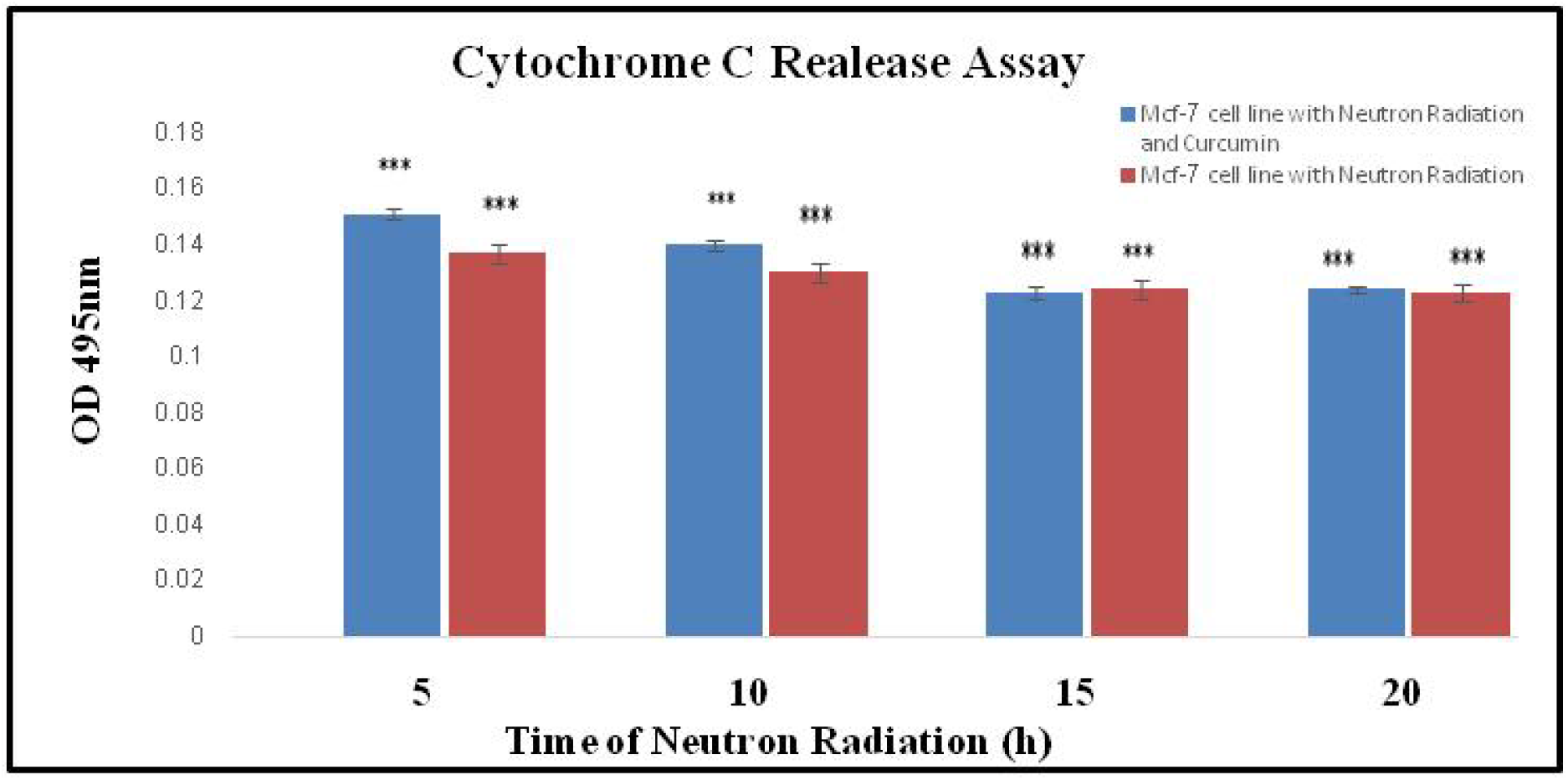
Measurement of cytochrome c released from mitochondria into cytosol at different times of neutron irradiation in the presence or absence of curcumin in MCF-7 cells in 3D culture Alkaline comet assay (***p<0.001)

#### 1.4.2.7 Alkaline comet assay

In this study, the alkaline comet was used to investigate the induction of cell death by neutrons on MCF-7 cells. The induction rate of neutron apoptosis at irradiation times of 5, 10, 15, and 20 h on MCF-7 cells in 3D culture was 5.5, 22, 24, and 26.5, respectively. This effect was equal to 6.5, 20.5, 22, and 23 in the presence of curcumin at the mentioned irradiation times (Figure 1-7). Statistical analysis showed that the effect of neutron radiation on the cell death induction in the presence and absence of curcumin was significant at all times except at 5 h. The results also showed that the effect of neutron radiation on cell death induction was independent of the presence or absence of the curcumin.

**Figure 1-7.**
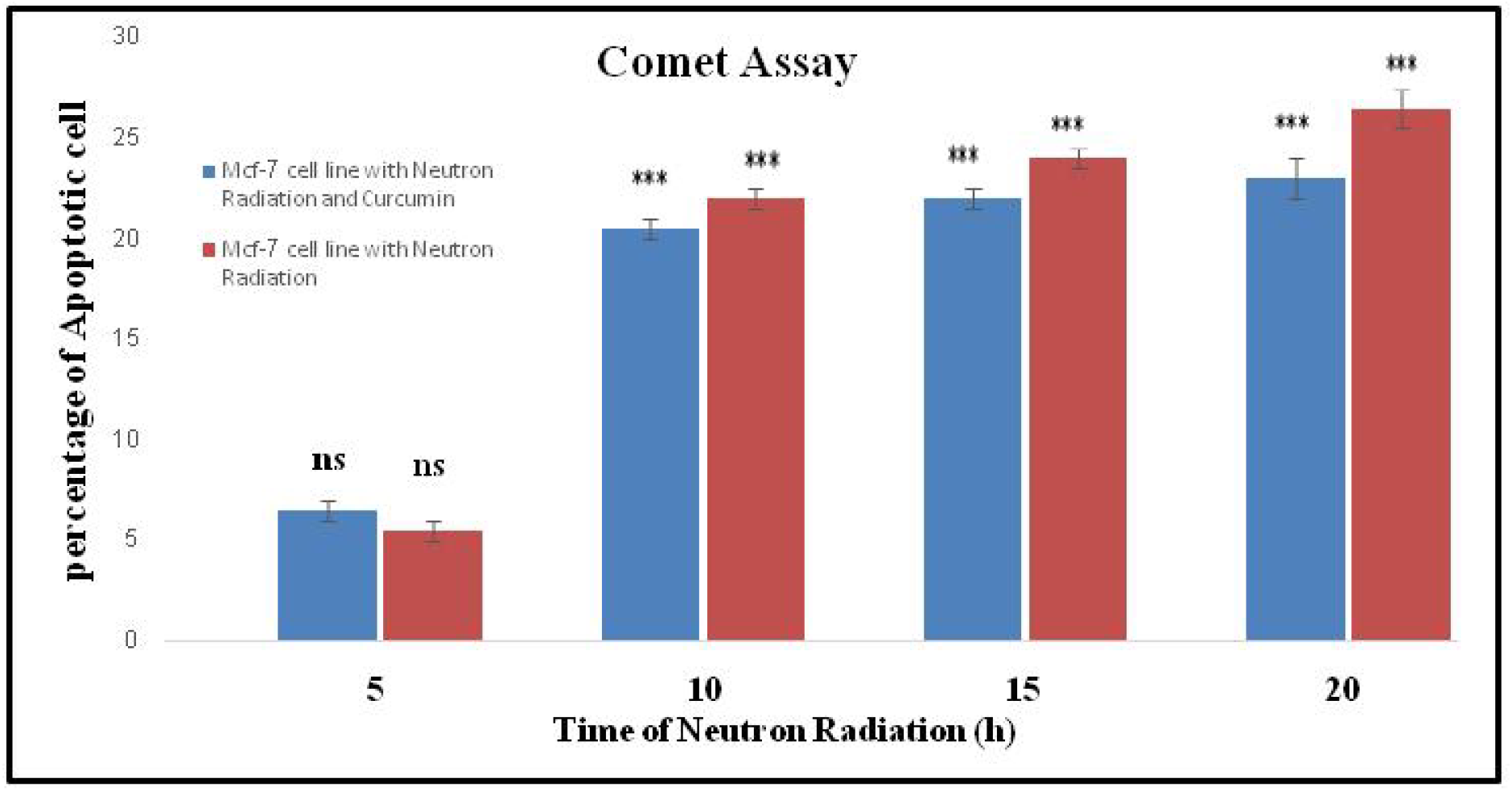
The rate of apoptosis induced at different times of neutron radiation in the presence and absence of curcumin in MCF-7 cells in 3D culture using Comet assay. (ns: not significant, ***p<0.001)

#### 1.4.2.8 Caspase-3 activity

The results showed that neutron radiation has a significant effect on the activity of caspase-3 enzyme in cells. Examination of irradiation times of 5, 10, 15, and 20 h on MCF-7 cells was 0.068, 0.071, 0.066, and 0.071, respectively. This effect of neutrons in the presence of the above times was equal to 0.066, 0.061, 0.068, and 0.063. The results show that the production of this enzyme in cells exposed to different times of neutron irradiation except at 5 hours compared to the control is significant (Figure 1-8) but the effect of neutron irradiation in the presence of curcumin and without it at the mentioned times is not significant.

**Figure 1-8.**
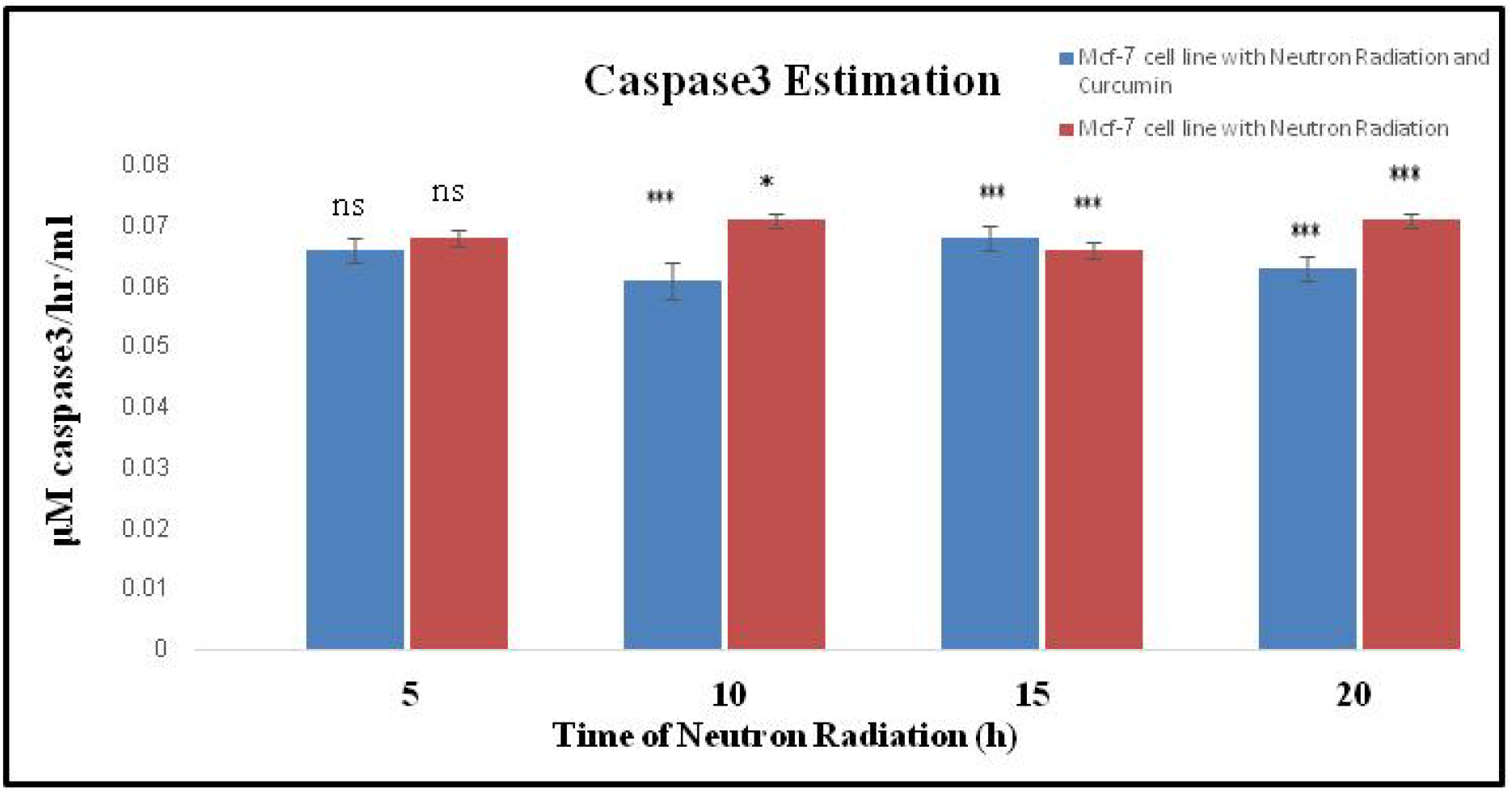
Caspase-3 activity in MCF-7 cells by examining the effect of neutrons in the presence and absence of curcumin using Caspase-3 activity assay. (ns: not significant, *p<0.05, ***p<0.001)

## 1.5. Discussion

Cancer is one of the major medical problems threatening human health today. Among cancers, the most common cancer among women is breast cancer (3). Reports indicate that treatments such as surgery, chemotherapy, and radiotherapy have played a limited but important role in the treatment of this disease. To increase the efficiency of treatment and increase the life of patients, new cancer treatment strategies based on the use of combined methods such as the use of conventional methods with intermediates of biological origin that have separate molecular mechanisms have been considered (6). One of the advantages of using these new methods is the reduction of systemic toxicity due to chemotherapy, radiation, or radiotherapy due to the reduction of the dose received by the patient (6). Curcumin in turmeric is one of the candidate molecules proposed in the treatment of cancer alone or combination with other methods. Result of recent studies have shown that this molecule has broad biological properties such as anti-inflammatory, antioxidant, anti-diabetic, and anti-cancer (8-7).

In this study, the effect of neutron radiation emitted from the ^241^Am-Be neutron source and the combined effect of neutrons and curcumin to increase the treatment efficiency of MCF-7 breast cancer in a 3D culture medium were considered. MTT, neutral red uptake assay, nitric oxide, glutathione assay, catalase, cytochrome c, comet assay, and caspase-3 were used to determine the effect and type of mortality due to neutron effect as well as the combined effect of neutron and curcumin in cancer cells. In this study, the MTT method was used to evaluate the cytotoxicity caused by the effect of neutron radiation as well as the combined effect of neutron radiation and curcumin (80 μM concentration) and the neutral red test was used to confirm the results. The results showed that neutron irradiation at 5, 10, 15, and 20 hours reduced the survival of tumor cells by 90, 65.96, 62.1, and 53.06%, respectively. However, the cell viability under the effect of neutron radiation and curcumin at these times was not statistically significant compared to the effect of neutron radiation alone and was 89.7%, 61.3, 64.9%, and 50.2%, respectively. The results showed that the rate of induction of apoptosis due to neutron effect at irradiation times of 5 to 20 hours, respectively, increased with increasing irradiation time. Statistical analysis showed that the rate of apoptosis due to the combined effect of neutrons and curcumin was not significant compared to the effect of neutrons. In 2011, Veeraraghavan et al. compared the effect of the combination of curcumin and radiation therapy on pancreatic cancer cells. They significantly achieved minimum survival, maximum mortality, and potential induction of apoptosis after combining 100 nM curcumin with radiation therapy and noted drug hypersensitivity in BxPC-3, Panc-1, and MiaPaCa-2 cell lines. On the other hand, their studies showed that the addition of curcumin before irradiation has better results in stopping the G2/m phase of the cell cycle than the addition of synchronous and radiotherapy (20). Zhan et al. (2014) also showed that curcumin increases the sensitivity of MCF-7 cells to chemotherapy drugs (21). Accordingly, the results have shown that curcumin can be a suitable candidate for use in combination cancer therapies as an intermediate molecule (22).

In 2004, Damondaran et al. Studied the effect of curcumin on prostate cancer cells and compared its results with the supplementation of curcumin with x-ray radiation therapy. The results showed that curcumin is a potent radiation sensitizer that inhibits the growth of prostate cancer cells and reduces cell survival factors as a result of radiation and increases the sensitivity of radiation in tumor cells. They showed that radiation therapy combined with curcumin compared to radiation alone could inhibit tumor growth and increase cell death in various types of cancer (23).

The neutron produced by the ^241^Am-Be source with 5-corie activity used in this study is a fast neutron and one of the most widely used neutron sources (24). Neutrons cause various interactions depending on their energy level when exposed to a living cell. Fast neutrons in contact with living tissue cells produce particles such as protons, alpha, and photons that have a destructive effect on DNA (25). This type of neutron damages the cell membrane and activates cell death pathways (24). The researchers’ findings show that neutrons are involved in hypoxia-induced cell death. This feature has led to the selection of neutrons to treat low-growth tumors such as prostate cancer (25). In general, neutrons are much more effective in destroying tumor cells than radiation with low linear energy transfer (LET), such as gamma, X, electron, etc. (26). The results of calculating the fast neutron flux around the source by two methods of simulation with MCNPX and detection with BF3, showed that with increasing the distance from the source, the Fast neutron flux decreased exponentially. According to the concordance of simulation results with MCNPX and detection with BF3 and based on cylindrical absorption dose measurements with vial/flask dimensions of cell culture (length 11 cm) at a distance of 22 cm, the maximum dose rate was calculated as 0.6 mGy per hour. The results of neutron dose rate measurement in the ^241^Am-Be source also showed that if the test cells in the vial are located at a distance of 22 cm from the inlet of the collimator and are exposed to neutron radiation for 5, 10, 15, and 20 hours, the dose The neutrons received by breast cancer cells will be 3, 6, 9 and 12 mGy per hour, respectively. It is noteworthy that the neutron dose emitted from the ^241^Am-Be source in less than 24 h, does not cause damage to normal living tissue cells (27).

Previous studies on the effect of curcumin on MCF-7 cell in 3D culture in alginate particles have shown that the IC50 of this substance was 80 μM (11). Zargan et al. (11) reported that this molecule reduced the survival of breast cancer cells by inducing apoptosis and necrosis, mainly apoptosis. In addition, the results of their study showed that curcumin reduces the production of cellular nitric oxide and increases the production of catalase and glutathione.

In this study, the effect of neutron radiation emitted from the ^241^Am-Be neutron source and the combined effect of neutrons and curcumin to increase the treatment efficiency of MCF-7 breast cancer in a 3D culture medium were considered. MTT, Neutral red uptake assay, Nitric oxide, Glutathione assay, Catalase, Cytochrome c, Comet assay, and Caspase-3 were used to determine the effect and type of mortality due to neutron effect as well as the combined effect of neutron and curcumin in cancer cells. In this study, the MTT method was used to evaluate the cytotoxicity caused by the effect of neutron radiation as well as the combined effect of neutron radiation and curcumin (80 μM concentration) and the neutral red test was used to confirm the results. The results showed that neutron irradiation at 5, 10, 15, and 20 hours reduced the survival of tumor cells by 90, 65.96, 62.1, and 53.06%, respectively. However, the cell viability under the effect of neutron radiation and curcumin at these times was not statistically significant compared to the effect of neutron radiation alone and was 89.7%, 61.3, 64.9%, and 50.2%, respectively. The results showed that the rate of induction of apoptosis due to neutron effect at irradiation times of 5 to 20 hours, respectively, increased with increasing irradiation time. Statistical analysis showed that the rate of apoptosis due to the combined effect of neutrons and curcumin was not significant compared to the effect of neutrons. In 2007, souto et al. investigated the effects of fast neutrons produced by the ^241^Am-Be source on polycarbonate materials and showed that fast neutrons at a dose rate of 0.5 to 20 mSv destroy carbon and oxygen bonds in materials, including living cells (28). In 2015, Saeed et al. investigated the effect of very low doses of fast neutrons (0.009 Gy) on the output of ^241^Am-Be with 5-curie activity *in vivo* on erythrocyte lipid membranes and proteins. Their result showed that the greatest effect of neutrons on cell membranes is due to their effect on methyl and methylene groups, which are composed of carbon and hydrogen (27). In 2016, Nafee et al. reported that neutrons alter or inactivate the protein responsible for cell transcription by breaking the c–o bond in RNA (29).

Examination of the molecular formula of curcumin (c_21_h_20_o_6_) shows that this molecule has a hydrocarbon structure and has multiple bonds c–o and c–h (30). According to the reports of Souto et al. (28), Saeed et al. (27), and Nafee et al. (29) It has been destroyed and the death of the breast cancer cell has been caused by neutron radiation. On the other hand, the induction of apoptosis in neutron-induced cancer cells in this study is consistent with a 2016 report by Nafee et al.

The results of neutron effect and combined effect of neutron and curcumin on the release of cytochrome c from mitochondria as well as activation of caspase-3 showed that the amount of cytochrome c release in the cytoplasm had an upward trend depending on the irradiation time and caspase-3 activation. The results of these two experiments were statistically significant compared to the control but not significant compared to each other. The results of this study is consistent with the results of Damondaran et al. (23).. The results of studying the effect of neutrons and the combination of neutrons and curcumin on the production of nitric oxide, catalase, and GSH also showed that their production in both cases was not statistically significant under the influence of different irradiation hours. The results of this study showed that curcumin reduces nitric oxide and increases the production of catalase and glutathione. In conclusion, the results of the present study showed that the neutron source ^241^Am-Be with the applied dose can destroy the c–o and c–h bonds of curcumin, resulting in virtual death and induced apoptosis in breast cancer cells, mainly via apoptosis. It was caused by neutron radiation. On the other hand, according to the results of Comet Assay and Caspase-3 experiments, although neutrons induced apoptosis in breast cancer cells, the death rate due to necrosis was much higher than apoptosis. Due to the significant anti-cancer effect of curcumin in 3D culture, the use of this molecule before or after neutron therapy is recommended.

